# Grazing and mowing enhance aquatic macroinvertebrate diversity of small artificial ponds in eutrophic landscape

**DOI:** 10.64898/2026.03.24.713891

**Authors:** Jana Petruželová, Jan Petružela, Alexandra Černá, Marie Kotasová Adámková

## Abstract

Artificial pond construction is widely used in wetland restoration, yet biodiversity outcomes depend strongly on design and subsequent management. We tested how different regimes (grazing, mowing, and no management) influence habitat structure, water conditions, and aquatic macroinvertebrate diversity in newly excavated experimental ponds within an eutrophic wetland in South Moravia (Czechia). Across four focal groups (Mollusca, Odonata, Coleoptera, Heteroptera), we observed rapid colonisation of the newly built ponds. Species richness and densities rose during early development, dropped after drying events, and then partially recovered, indicating repeated “resetting” of communities under fluctuating hydrology. Periodic drying also prevented fish stock establishment. Management significantly affected species composition and both grazed and mowed ponds displayed higher densities (abundances) than controls, but differed only slightly in terms of species richness. The grazed ponds were characterised by high sunlight exposure, reduced reed dominance, and trampling-generated high littoral heterogeneity. These ponds showed highest numbers of taxa adapted to shallow and warm waterbodies, muddy substrate, semiaquatic microhabitats, or newly emerged and disturbed habitats. The mowed ponds promoted dense submergent vegetation, supporting Odonata representation and other taxa requiring aquatic vegetation. The control ponds remained highly shaded by high-grown reed, organic-matter rich, hosting a set of taxa tolerant of low-light, low-oxygen conditions. At the wetland scale, multiple small ponds increased overall diversity through high between-pond heterogeneity. Our results highlight that pond construction alone is insufficient for wetland restoration: follow-up long-term management regimes, especially extensive grazing, can rapidly generate structural heterogeneity and sustain diverse aquatic invertebrate assemblages in eutrophic wetlands.

## 1. Introduction

Wetlands are among the most important and threatened ecosystems on Earth (Maltby & Barker 2009, Kingsford et al. 2016, Tickner et al. 2020), providing a wide range of ecosystem services, including carbon sequestration, flood detention, water storage and nutrient retention (Vymazal 2010, Verhoeven 2014, Mitsch & Gosselink 2015). These biotopes are also biodiversity hotspots for many taxa, including plants, invertebrates and vertebrates, especially amphibians and birds, for which these habitats serve as important migratory and nesting grounds (Davies et al. 2008, Verhoeven 2014). However, a significant proportion of wetlands has been lost globally. In Europe, it is estimated that 50–80% have disappeared, with many others remaining degraded (Fluet-Chouinard et al. 2023). Alluvial wetlands have been particularly impacted due to the expansion of agricultural land and urban areas, eutrophication, pollution, the proliferation of invasive species, and the regulation of rivers (Sánchez-Carrillo et al. 2010, Altieri et al. 2022, Datta et al. 2022, Fluet-Chouinard et al. 2023). Therefore, these biotopes are extremely important for biodiversity conservation and restoration efforts (Davies et al. 2008, Verhoeven 2014, Tickner et al. 2020, Erwin 2009).

The creation of artificial ponds is becoming an increasingly widespread method of wetland restoration (Drayer & Richter 2016, Oertli 2018, Zhang et al. 2020, Moor et al. 2024, Fahy et al. 2025). Building wetland pondscapes has gained popularity in Czechia: over the past two decades, many such habitats have been created either on arable land or within degraded or even natural wetlands in order to increase local biodiversity (Devánová et al. 2026, Janáč et al. 2025). However, this approach is facing significant challenges. Artificial ponds often have monotonous shorelines that are unsuitable for diverse aquatic vegetation (Law et al. 2019, Zamora-Marín et al. 2021), which consequently reduces habitat offer for aquatic invertebrates (Thornhill et al. 2018). Additionally, larger ponds are colonised by non-native fish species, adversely affecting water and habitat quality (Janáč et al. 2025). Consequently, invertebrate and amphibian diversity in artificially created ponds is often significantly lower than in the natural ones (Pearl et al. 2005, Schulse et al. 2010, Denton & Richter 2013, Zamora-Marín et al. 2021). Finally, many artificial ponds lack a long-term management strategy, which is essential for sustaining them in a favourable state, especially in eutrophic and highly modified landscapes (Oertli 2018, Zamora-Marín et al. 2021).

There are several regimes for systemic wetland management, with interventions including grazing, mowing, burning and others as methods for invasive and expansive vegetation control (van der Walk 2009, Lamers et al. 2015). In particular, extensive livestock grazing is an increasingly important practice, as it has traditionally been used to utilise wetlands in many parts of Europe (van Diggelen et al. 2006, Biró et al. 2019). Studies have reported positive impacts of grazing on wetland diversity (Maltby & Barker 2009, Morris & Reich 2013, Marty 2015), including that of aquatic invertebrates (Marty 2005, Davis & Bidwell, 2008), although other reports found no benefits or even deleterious effects in the case of intensive pasture (Steinman et al. 2003, Bloechl et al. 2010, Epele & Miserendino 2015). Nevertheless, these studies consider wetlands in general, and the effects of management on newly constructed artificial ponds remain largely unexplored. This is unfortunate, as these interventions have high potential to improve pond condition. Biomass removal reduces vegetation closure and increases light availability within ponds (van der Walk 2009). Additionally, grazing in particular introduces a continuous, spatially uneven disturbance regime: trampling alters microtopography, producing shallow depressions and variable depth profiles (Reeves & Champion 2004, Middleton 2016, Fløjgaard et al. 2022). These processes increase small-scale habitat heterogeneity, which is widely recognised as a key driver of biodiversity (Whittaker & Levin 1977, Pickett & White 1985). Therefore, investigating the impact of continuous management on newly constructed ponds is particularly relevant, as it directly addresses the usual limitations of such artificial systems: expansive vegetation proliferation and structural homogeneity.

In this study, we selected a degraded alluvial wetland located within Czechia, designated for restoration through grazing and mowing combined with the creation of small artificial ponds (Kotasová Adámková et al. 2022). We conducted a field experiment to test the effects of different management treatments (grazing, mowing, and no management as a control) on aquatic macroinvertebrate diversity and on the characteristics of newly created ponds. Specifically, we tested whether the treatments significantly affect (i) species richness, abundance, and species composition of selected macroinvertebrate groups, and (ii) physico-chemical and habitat characteristics of the ponds. Additionally, we assessed the development and succession of the experimental ponds over time.

## 2. Materials and methods

### 2.1 Study area

The experimental area is a lowland wetland of 11 ha area situated on alluvial deposits of the *Spálený potok* stream, located in the municipality of Krumvíř in the south-eastern part of Czechia (South Moravia region), at an altitude of 179 m (48.9977197N, 16.9071019E). Until the 1970s, it had the character of a periodically flooded wetland meadow with halophilic vegetation (Kotasová Adámková et al. 2022), much of which was used for communal pasture and hay harvesting. Since the 1970s, the traditional farming there was abandoned, and the area began to face strong eutrophication due to increased phosphorus and nitrogen inputs from the *Spálený potok* watershed and the absence of management. Consequently, most of the wetland was overgrown by expansive common reed (*Phragmites australis* (Cav.) Trin. ex Steud.) at the turn of the 20^th^ and 21^st^ centuries (Fig. S1). In 2003, a pond approximately 1 ha in size and about 2 metres deep was constructed in the western part of the wetland (in this study, referred to as the *old pond*; Fig. 1, Fig. S1). For several years, it was used for recreational fishing, however, without a stable fish stock. Due to high nutrient input, its permanent character, and unfavourable morphology (i.e. great depth, almost vertical banks) it has been subjected to a dynamic succession. The water column has become heavily overgrown by submergent vegetation, and its dead biomass is gradually reducing the pond’s depth.

**Fig. 1.**
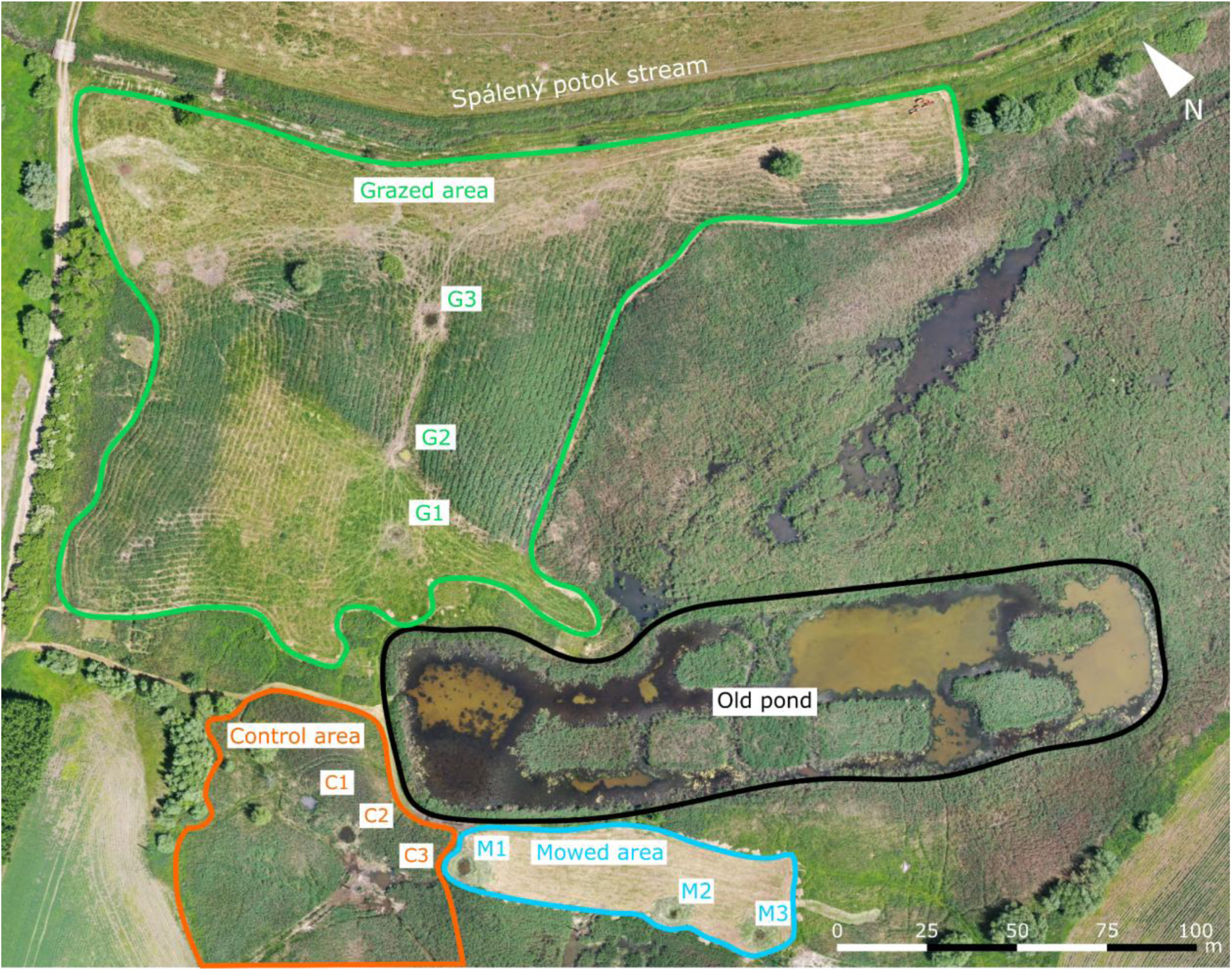
Aerial image of the study site taken on 22 June 2022. Polygons indicate managed areas (mowed and grazed), control area and the old pond. Codes M1–3, G1–3 and C1–3 denote the experimental ponds. The image was acquired using an unmanned quadcopter (registration number OK-X071A). Photo by L. Tichý.

In 2020, management in the form of grazing and mowing was initiated in the western part of the wetland (Fig. 1, Fig. S1, Fig. S2), and nine small experimental ponds were excavated to support local biodiversity (Marie Kotasová Adámková et al. 2022). The experimental ponds were built in September 2020, each with approximately the same size (circular shape with a 5 m diameter) and a maximum depth of about 1 m. Three ponds are located in the grazed area (G1–G3), three in the mowed area (M1–M3), and three in the control area (i.e. area without management; C1–C3; Fig. 1). The soil from the excavation was spread around the ponds. The experimental ponds partially dry up each summer; in dry years, some of them dry out completely for several weeks. Conversely, during heavy precipitation events, they are flooded for several days along with the nearby area. The strong water level fluctuations do not allow permanent survival of fish in the ponds (some individuals appeared in several ponds for a few weeks).

Due to the high productivity of the pasture, it is managed under an extensive grazing regime with a stocking rate between 1 and 2 livestock units per hectare. Grazing is carried out from late April–early May to late October–early November and residual vegetation is mechanically mowed and removed during winter. Grazing is performed by a combination of Aberdeen Angus and Aubrac cattle, representing heavier and lighter beef breeds. The cattle have unlimited access to ponds G1–G3, using them as a drinking supply.

The mowed area is cut twice annually using light machinery, including the littoral emergent vegetation of ponds M1–M3. All biomass is carefully removed by manual raking. A portion of the harvested biomass is used as cattle fodder, while the remainder is deposited in less conservation-valuable parts of the site to create microhabitats for reptiles. The control ponds (C1–C3) are left unmanaged and are characterized by dense, tall reed stands.

### 2.2 Aquatic macroinvertebrates sampling

The data on aquatic macroinvertebrates were obtained from nine experimental ponds and the old pond. Samplings were performed each September and June in the period between September 2020 (two weeks after the experimental ponds were built) and June 2024 (i.e., 8 samplings in total). The old pond was additionally sampled also in June 2020 in pre-management period. Macroinvertebrates were collected by a semiquantitative method using a hand net with frame 25×25 cm and mesh size 0.5 mm in the littoral zone of each waterbody. In each experimental pond, 20 patches of approximately 25×25 were sampled, or ten patches in case of low water level in dry periods (as 20 patches would be very invasive for very small waterbodies). In the old pond, 20 patches were always sampled. The sampling was performed to proportionally capture all mesohabitats available in each waterbody (submergent macrophytes, reed, organic substrate, mud etc.). Each sample was placed on a white plate and examined directly in the field. Invertebrates were transferred with tweezers to sample containers with fixing mediums (individuals of easily recognizable species were identified in the field and released). Four macroinvertebrate groups dominant in these ponds were selected: Mollusca, Odonata (larvae), and Coleoptera and Heteroptera (adults and larvae). Other groups were omitted due to very low numbers of species (Ephemeroptera), rare occurrence (Trichoptera), or difficult determination (Diptera, Annelida). Fixed individuals were transferred to the laboratory and determined by specialists to the lowest possible level (mainly species). The information about the macroinvertebrate species ecology stated in the text was sourced from the following literature: Mollusca (Welter-Schultes 2012); Odonata (Boudot & Kalkman 2015); Heteroptera (Tempelman & van Haaren 2009); Coleoptera (Boukal et al. 2007; Klausnitzer 2009; Boukal et al. 2012). The raw species records data are available at Mendeley Data repository (Petruželová et al. 2026)

### 2.3 Physico-chemical and habitat variables

During each sampling term and in each waterbody, physico-chemical variables and habitat variables were obtained in the field, and water samples were collected for laboratory analyses. Water temperature, dissolved oxygen, oxygen saturation, pH and conductivity were measured in situ using a portable instrument (HACH HQ40d; HACH Co., Loveland, CO, USA). Water transparency was measured by Secchi disc, maximum depth was measured using a meter. Water samples were filtered and analysed in the laboratory within 24 h of sampling for concentrations of total phosphorus (TP), total nitrogen (TN) and total organic carbon (TOC). Habitat characteristics in the littoral zone were visually estimated in the field. These include percentage cover by filamentous algae and different types of macrophytes: submergent (in the old pond *Ceratophyllum demersum* L., in experimental ponds predominantly *Chara vulgaris* L. and *Ch. hispida* L.), reed (*Phragmites australis*), “other emergent macrophytes” (e.g. *Carex* sp., *Ranunculus sceleratus* L., *Mentha* sp.), and natant macrophytes (*Lemna minor* L.). Lastly, we estimated the particulate organic matter (POM) cover, shading (i.e. proportion of littoral shaded be high-grown reed) and littoral slope (ordinal variable with levels 1–5 according to prevailing character of the banks: 1 – extremely sloping, almost vertical banks; 2 – strongly sloping; 3 – medium sloping; 4 – low sloping; 5 – very low sloping, with high proportion of aquatic-terrestrial habitat transitions). The raw environmental and physico-chemical variables data are available at Mendeley Data repository (Petruželová et al. 2026)

### 2.4 Data analysis

Total taxa richness and macroinvertebrate group richness were recorded for each sampling period and pond. Larval and adult taxa in Coleoptera (and certain Heteroptera) were kept separately, because (i) many larval taxa cannot be adjusted to adult taxa, as they are usually determined to a higher taxonomic level, and (ii) larvae often have different ecology from adults (in Coleoptera). As individual samples were taken from 20 or 10 patches, all taxa abundances were recalculated as densities, i.e. the number of individuals per unit of sampled area of 1.25 m^2^ (which corresponds to 20 sampling patches of 25×25 cm). Densities were log-transformed and used in the analyses of community composition. The Spearman correlation table (Spearman 1904) was calculated to examine correlations within variables. Variable *dissolved oxygen* was excluded from analyses for high correlation with *oxygen saturation* (ρ = 0.98).

To assess temporal trends in species richness and abundance within the experimental ponds, we analysed the data using generalised additive models with mixed effects (GAMM) using the “mgcv” package in R (Wood 2017). Both response variables were modelled as overdispersed count data with a negative binomial error distribution and a log link. Sampling time was included as a categorical fixed effect to estimate differences among sampling terms without imposing a smooth or monotonic temporal trend. Management regime was included as a fixed effect, and pond identity was included as a random intercept to account for repeated sampling of the same ponds. Model parameters and smoothing penalties were estimated using restricted maximum likelihood (REML). Identical model structures were used for both abundance and species richness. Additionally, we assessed temporal trends in species richness and abundance in the old pond using Spearman’s ρ (Spearman 1904).

To assess the significance of management regime for species richness, abundances, and environmental variables, we used rank-based nonparametric longitudinal analysis (nparLD) with an F1-LD-F1 design (Brunner et al. 2002), to account for repeated sampling of the same ponds over time. Pond identity was specified as the subject (repeated-measures) factor, management regime (C/G/M) as the between-subjects factor, and sampling time as the within-subjects factor. When a regime effect was detected, we performed post-hoc pairwise comparisons among regimes using the nparLD pairwise procedure. To account for multiple testing across response variables, we adjusted regime-effect p-values using the Benjamini–Hochberg FDR. The analysis was conducted using the nparLD 2.2 R-package (Noguchi et al. 2012).

Distance-based Redundancy Analyses (db-RDA) on Bray-Curtis distances of the species composition data were performed (i) using sampling site as a conditional variable and sampling term (*Time*) as an explanatory variable to display temporal changes, and (ii) using management regime as an explanatory variable to evaluate the impact of different management regimes on the experimental ponds community composition (the first sampling term was excluded from this analysis due to outlying low diversity). The significance of the effects of explanatory variables was tested using a 9999-permutation procedure, and the adjusted R^2^ was used as an unbiased estimate of explained variation (Peres-Neto et al., 2006). Variables that differed significantly between the management regimes were post-hoc fitted into the second ordination diagram (function “envfit” in “vegan” package; Oksanen et al. 2025). Additionally, we computed the Bray-Curtis distances between the three management regimes for each sampling term to visualise the temporal development of between-regime community heterogeneity. Indicator analysis with 1,000 randomisations was applied to log-transformed density data to identify indicator taxa for the three types of experimental pools (“indval” function from the “labdsv” package, Roberts 2025).

## 3. Results

### 3.1 Diversity, species richness and abundance: general patterns and development in time

In total, 145 taxa of macroinvertebrates from the four selected groups were recorded in the study period (see list of all taxa in Table S1) in all studied ponds (experimental ponds and the old pond). Most of them represent Coleoptera (96 taxa), followed by Heteroptera (25), Odonata (14) and Mollusca (10). Most of the taxa (93) were found both in the old pond and in the experimental ponds (at least once), 39 taxa were unique to the experimental ponds, and 13 taxa were found only in the old pond.

The total species richness in experimental ponds showed a statistically significant trend (GAMM, p < 0.001, deviance explained = 68.9 %) over the study period (Fig. 2A). All sampling times showed significantly higher richness than the reference term (September 2020), except for September 2022 (p = 0.135). Richness increased in most ponds from the start of the sampling until June 2022, when it peaked (maximum of 36 taxa). In the next sampling in September 2022 (after a dry summer, when all experimental ponds dried completely or significantly), species richness dropped and then only slightly increased until the end of the study period (Fig. 2A). Strong variability among individual ponds was observed within a given management regime (p < 0.001) but differences among management regimes were negligible. Density showed a similar trend as total species richness (Fig. 2B; GAMM, p < 0.001, deviance explained = 77.2 %), again with all sampling times significantly higher than the reference with the exception of September 2022 (p = 0.323). In contrast to the species richness, the management regime had a significant effect, with grazing displaying the significantly higher densities than control (p < 0.001) on the density trend, with the additional pond identity effect again being significant (p = 0.005). In the old pond, we found a weakly significant decreasing temporal trend for densities (ρ = **−**0.68, p = 0.042), but no significant temporal trend for species richness (ρ = **−**0.48, p = 0.194; Fig. 2).

**Fig. 2.**
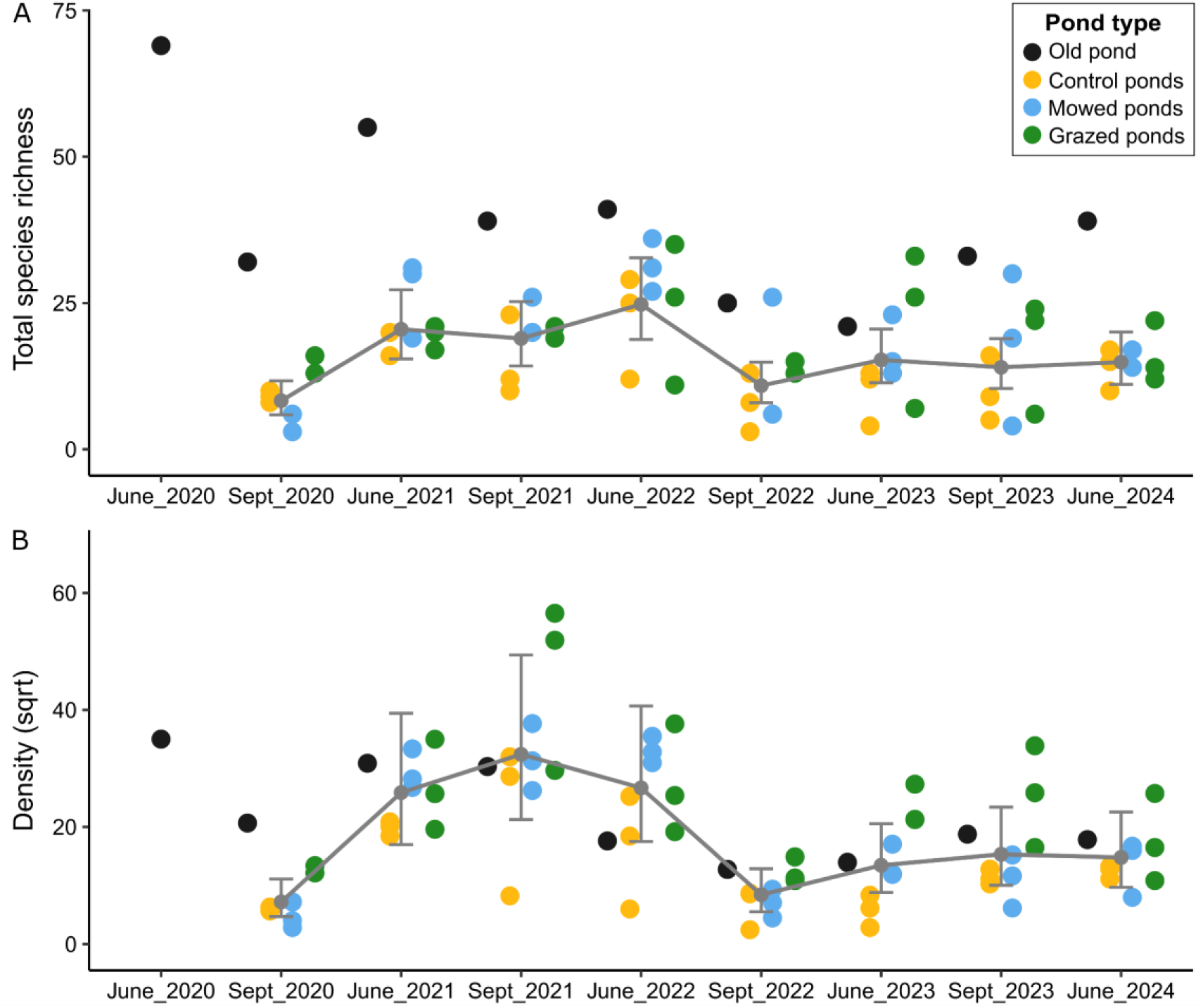
Development of species richness (A) and density (B) of macroinvertebrates in ponds over time. The type of aquatic biotope is indicated by colour. In June 2020, only the old pond was sampled; first sampling of the experimental ponds was performed two weeks after they were built in September 2020. The grey curves represent the modelled temporal dynamics of the experimental ponds from the GAMM regression (the old pond data did not enter the regression modelling).

The development of species richness differed between the four studied groups of macroinvertebrates (Fig. S3). Mollusca first appeared in the experimental ponds during the second sampling, and the number of species then remained constant between one and four species per pond and sampling time (except few outliers). Odonata larvae also appeared in the experimental ponds in the second sampling, sharply increasing up to 7 species per pond. But, in the second half of the study period, only zero to three taxa were found at each pond (Fig. S3). Heteroptera and Coleoptera appeared in the ponds during the first sampling period (adults taxa only), and their species richness trends over time followed the trend of total species richness. Heteroptera gained numbers up to 11 species, and Coleoptera up to 22. Densities of individual macroinvertebrates groups showed similar trends as the total density (Fig. S3).

The old pond density values at most samplings ranged around the average values reached by the experimental ponds (Fig. 2B). Its species richness fluctuated strongly in time (mean = 39; Table 1) and was only a few times exceeded by an individual experimental pond richness (Fig. 2A). However, the total (gamma) diversity of all the exp. ponds was, except the first sampling, always higher (mean = 55; Table 1). A large difference between gamma and alpha diversity (mean = 17) of exp. ponds led to their high beta diversity, which was moreover gradually increasing through our study period (Table 1). Diversity of the wetland (i.e. total number of species in the old pond and experimental ponds) ranged from 47 to 91 species at each sampling (mean = 68; Table 1).

**Table 1.** Diversity indices in the studied ponds based on macroinvertebrate data for each sampling. α – alpha diversity (mean species richness) of the experimental ponds; γ – gamma diversity (total richness) of all experimental ponds; β – beta diversity of the experimental ponds; Old P – species richness of the old pond; Wet – wetland species richness (i.e. total richness of all experimental ponds and the old pond).

| Sampling | $\alpha$ | $\gamma$ | $\beta$ | Old P | Wet |
| --- | --- | --- | --- | --- | --- |
| June 2020 | - | - | - | 69 | - |
| Sept 2020 | 8.9 | 23 | 2.59 | 32 | 47 |
| June 2021 | 21.1 | 64 | 3.03 | 55 | 78 |
| Sept 2021 | 19.6 | 51 | 2.61 | 39 | 60 |
| June 2022 | 25.8 | 82 | 3.18 | 41 | 91 |
| Sept 2022 | 11.4 | 43 | 3.76 | 25 | 60 |
| June 2023 | 16.2 | 61 | 3.76 | 21 | 67 |
| Sept 2023 | 15.0 | 57 | 3.80 | 33 | 70 |
| June 2024 | 15.3 | 59 | 3.85 | 39 | 74 |

### 3.2 Species richness and abundance: impact of management on the experimental ponds

Total species richness of macroinvertebrates did not significantly differ between the types of experimental ponds (Table 2, Fig. 3). Considering individual groups of macroinvertebrates, only Odonata species richness showed a difference, as it was higher in the mowed and marginally also in the grazed ponds in comparison with the control ponds. In Mollusca and Coleoptera the numbers of species were often higher in the managed ponds, but the differences were not significant. In Heteroptera, the richness was comparable between the types. The total density in grazed ponds was significantly higher than in the control ponds, and in the mowed ponds there was a marginally insignificant difference (Table 2, Fig. 3). In Odonata, all three types of experimental ponds differed in densities, with the highest values in the mowed, medium in the grazed, and the lowest in the control ponds. The Mollusca and Coleoptera densities were higher in the grazed ponds in comparison with the other two types. Densities of Heteroptera were comparable between all types of experimental ponds.

**Fig. 3.**
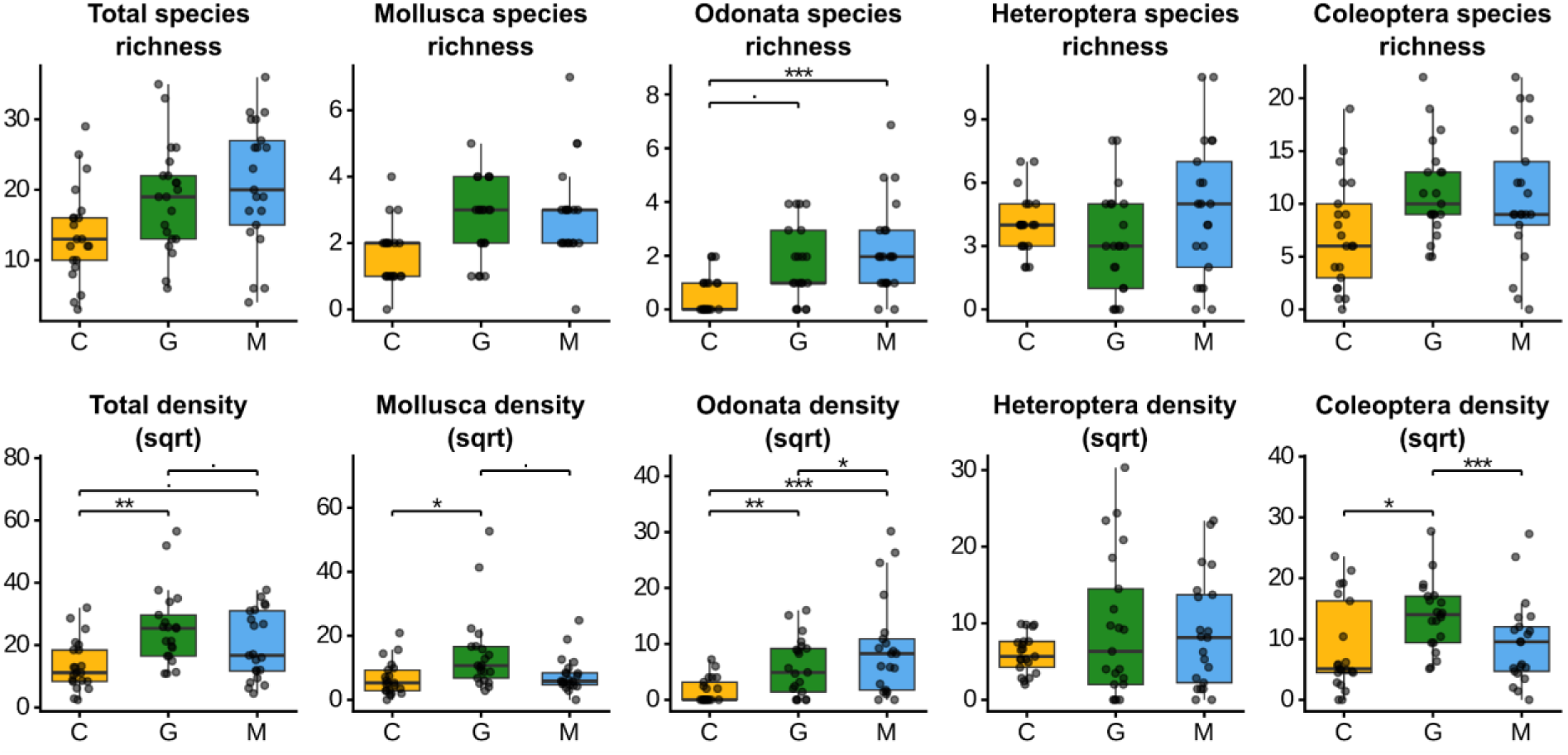
Species richness and density in individual types of experimental ponds (C – control, G – grazed, M – mowed). Data since June 2021 were used (the first sampling was omitted due to outlying low diversity). Boxes represent the median (horizontal line) and inter-quartile range (25^th^–75^th^ percentiles) and whiskers represent the range of values within 1.5 box-lengths. Points along boxplots stand for individual values. Signs above plots indicate significant differences between connected boxes (see Table 2). *** p < 0.001; ** p < 0.01; * p < 0.05; . p < 0.

**Table 2.** Summary of the nonparametric longitudinal model with post-hoc pairwise comparisons with FDR corrected p-values performed on species richness and abundance in the three types of experimental ponds. Data since June 2021 were used (the first sampling was omitted due to low initial diversity). Parameters are ordered from the most significant to the least significant. For graphical visualisation of the results, see Fig. 3. p (raw) – uncorrected p-values from the model; p (FDR) – FDR corrected p-values from the model; C vs. G, C vs. M, G vs. M – FDR corrected p-values and significance from post-hoc pairwise comparisons of the experimental ponds (C – control, G – grazed, M – mowed). *** p < 0.001; ** p < 0.01; * p < 0.05; . p < 0.1.

| Variable | p (raw) | p (FDR) | C vs. G | C vs. M | G vs. M |
| --- | --- | --- | --- | --- | --- |
| Odonata density | <0.001 *** | 0.001 *** | 0.007 ** | <0.001 *** | 0.032 * |
| Odonata sp. richness | 0.003 ** | 0.013 * | 0.056 . | <0.001 *** | 0.272 |
| Total density | 0.004 ** | 0.013 * | 0.005 ** | 0.09 . | 0.095 . |
| Coleoptera density | 0.016 * | 0.04 * | 0.017 * | 0.734 | <0.001 *** |
| Mollusca density | 0.034 * | 0.068 . | 0.012 * | 0.533 | 0.085 . |
| Mollusca sp. rich. | 0.176 | 0.257 | - | - | - |
| Coleoptera sp. rich. | 0.18 | 0.257 | - | - | - |
| Total sp. richness | 0.261 | 0.328 | - | - | - |
| Heteroptera sp. rich. | 0.567 | 0.631 | - | - | - |
| Heteroptera density | 0.921 | 0.921 | - | - | - |

### 3.3 Species composition and ponds characteristics: influence of management and time

Species composition of macroinvertebrates in ponds was significantly affected by both time and management (Fig. 4). Time explained 7.01% of variation in the entire dataset (db-RDA, p < 0.001). While most of the experimental ponds communities developed along the first ordination axis constrained by time, the old pond does not show visible temporal development, with individual samplings circling along the center of the ordination diagram (Fig. 4A). The changes in species composition of the exp. ponds are connected to changes in species richness (Fig. 2A), relative abundances of the taxa, and taxa exchange (Table S1) throughout the study period. The most distinct group of sites consists of the samples of the first sampling term in September 2020 (upper left corner in Fig. 4A), when the communities were composed of very few species and individuals (Fig. 2). For this distinctness, the first sampling term of the exp. ponds was excluded when testing the impact of management. Management type explained 7.24% of variation in macroinvertebrate species composition in the experimental ponds (db-RDA, p < 0.001; Fig. 4B). The grazed ponds differed from the control ponds most significantly (G vs. C: p < 0.001, 8.3%), while differences between the mowed and control ponds and the mowed and grazed ponds were lower (M vs. C: p < 0.002, 3.5 %; M vs. G: p < 0.002, 4.8 %).

**Fig. 4.**
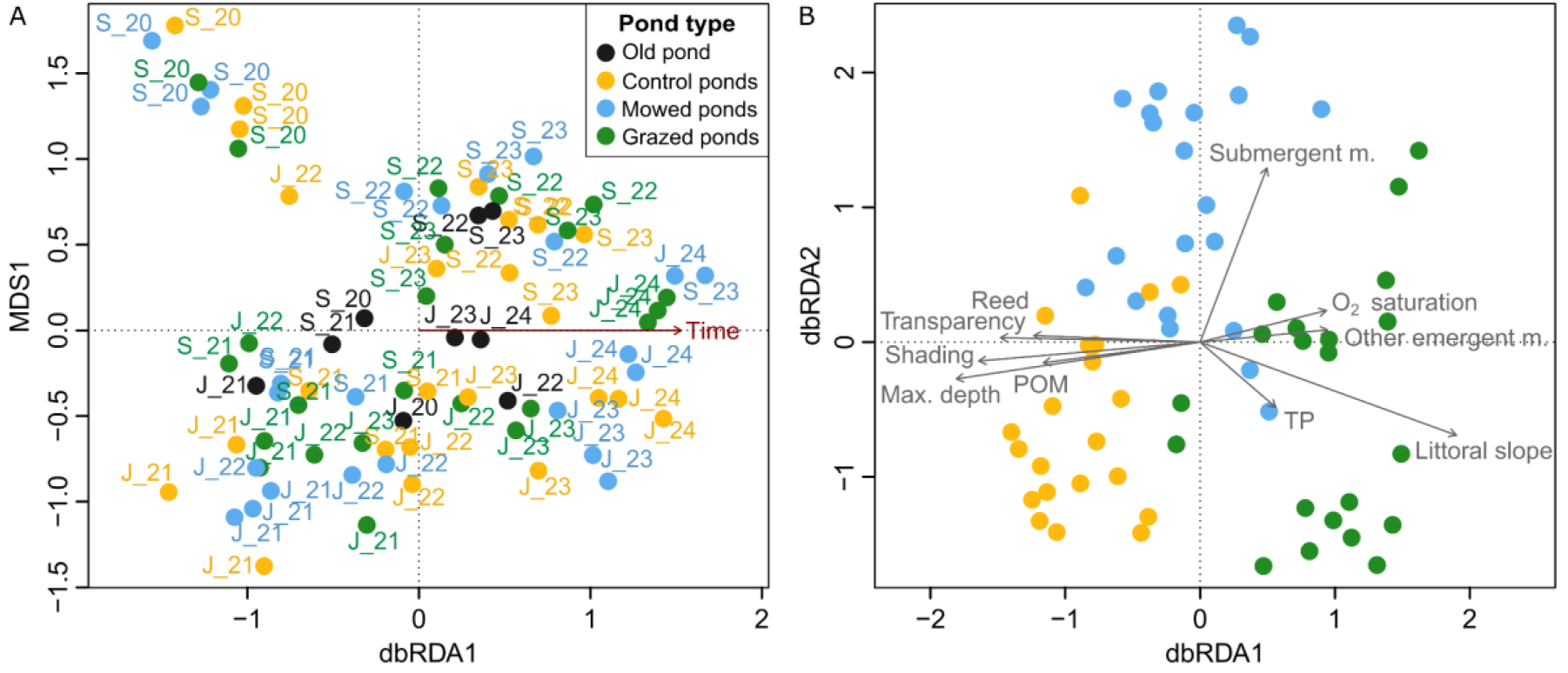
Ordination plots of db-RDA analyses on macroinvertebrate community composition of the ponds: (A) db-RDA with time used as the constraining variable (first axis; explaining 7.01% of variation) on the entire dataset, (B) db-RDA on the experimental ponds dataset, excluding the old pond data and the first sampling term (Sept. 2020), using the management as the only constraining variable (explaining 7.24% of variation). Environmental variables significantly differing at least in one pair of the experimental pond types (see Table 4) were passively fitted into the ordination and shown as arrows.

The difference (Bray-Curtis dissimilarity) between the types of experimental ponds showed a specific trend in time (Fig. 5). Initial species poor communities in September 2020 differed moderately (approx. 0.48–0.6), then the dissimilarities dropped for three sampling periods to ∼0.38 in the mowed vs. grazed ponds and 0.4–0.48 in the mowed vs. control ponds, while the grazed vs. control ponds dissimilarities remained about 0.5. Since the ponds refilled after dry summer, all dissimilarities have increased since September 2022 (but with a certain drop in September 2023), with the grazed vs. control ponds gaining the highest differences (up to 0.66). Across time, the pair showing the highest dissimilarity tend to be the grazed vs. control ponds, followed by the control vs. mowed ponds. The grazed and mowed ponds incline to have the most similar communities across time (Fig. 5).

**Fig. 5.**
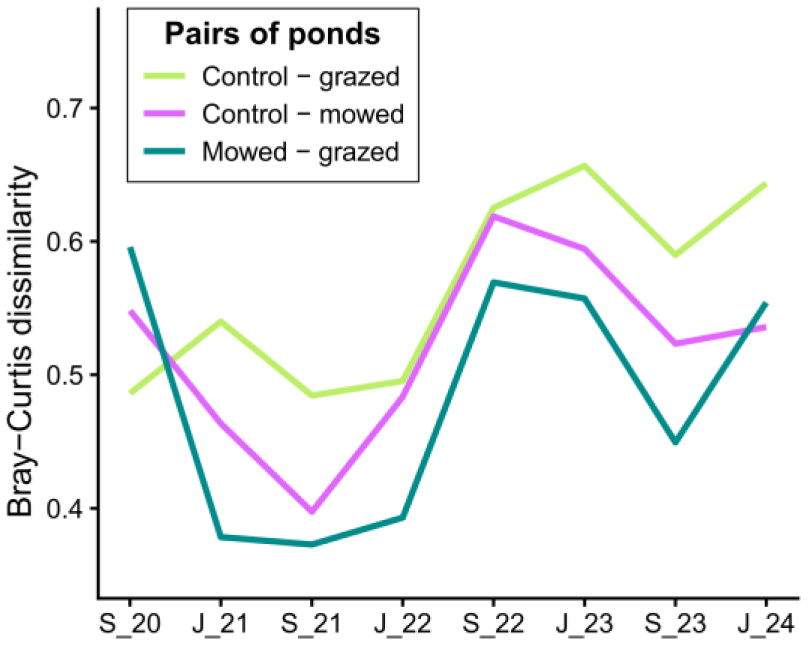
Time development of Bray-Curtis dissimilarity values between pairs of types of experimental ponds, counted on log-transformed species composition data. X axis stands for sampling terms between June 2020 and June 2024 (J – June, S – September, 20 – 2020, 21 – 2021, 22 – 2022, 23 – 2023, 24 – 2024).

The community composition of the three types of experimental ponds can be characterised by indicator taxa (Table 3). The highest number of indicator taxa was found in the grazed ponds (9) comprising mostly species (adult taxa) of Coleoptera (7), one species of Mollusca and one of Heteroptera. In the mowed ponds, five taxa were significant, counted among Heteroptera (2), Mollusca (2) and Odonata (1). Only two taxa, one representative of Coleoptera and one of Mollusca, were identified as indicator taxa in the control ponds.

**Table 3.** Results of the indicator analysis, performed on species-composition data of the experimental ponds.

| Taxon | Group | Type | Indicator value | p-value |  |
| --- | --- | --- | --- | --- | --- |
| *Aplexa hypnorum* | Mollusca | Control | 0.308 | 0.018 | * |
| *Contacyphon* sp. lv. | Coleoptera | Control | 0.221 | 0.032 | * |
| *Gyraulus crista* | Mollusca | Mowed | 0.410 | 0.001 | ** |
| *Ilyocoris cimicoides* | Heteroptera | Mowed | 0.363 | 0.001 | ** |
| *Plea cryptica* | Heteroptera | Mowed | 0.360 | 0.025 | * |
| *Sympetrum striolatum* | Odonata | Mowed | 0.287 | 0.022 | * |
| *Lymnaea stagnalis* | Mollusca | Mowed | 0.251 | 0.048 | * |
| *Radix labiata* | Mollusca | Grazed | 0.640 | 0.001 | ** |
| *Hygrotus impressopunctatus* | Coleoptera | Grazed | 0.451 | 0.001 | ** |
| *Hydroglyphus geminus* | Coleoptera | Grazed | 0.448 | 0.001 | ** |
| *Agabus bipustulatus* | Coleoptera | Grazed | 0.345 | 0.008 | ** |
| *Helophorus* gr. *minutus* | Coleoptera | Grazed | 0.318 | 0.003 | ** |
| *Ochthebius bernhardi* | Coleoptera | Grazed | 0.290 | 0.003 | ** |
| *Ochthebius minimus* | Coleoptera | Grazed | 0.286 | 0.003 | ** |
| *Sigara nigrolineata* | Heteroptera | Grazed | 0.244 | 0.012 | * |
| *Enochrus quadripunctatus* | Coleoptera | Grazed | 0.169 | 0.033 | * |

The experimental ponds significantly differed also in terms of physico-chemical and habitat variables (Table 4, Fig. 4B, Fig. 6). The most distinct pair of ponds was the control and grazed ponds, both in terms of number of differing variables and significance of the differences, while the least distinct were the grazed and mowed ponds (Table 4). The most significant variables were those describing the habitat structure of the ponds. Littoral slope was usually medium in both mowed and control ponds, but very low in the grazed ponds for most of the study period (Fig. 6, Fig. S4). Shading was minimal most of the time in the grazed ponds, high in the control ponds, and medium in the mowed ponds (Fig. 6). There was high dispersion of values, depending on season and management: lower shading was at the beginning of the study period, in June samplings, and in the mowed ponds also shortly after mowing (Fig. S4). POM cover was highest in the control ponds most of the time, medium in the mowed and lowest in the grazed ponds (Fig. 6). POM fluctuated with seasons, after dry periods it mineralised and decreased (Fig. S4). Oxygen saturation and other emergent macrophytes cover were lowest in the control ponds (Fig. 6). Water transparency was lowest in the grazed ponds, but it has to be mentioned that this variable was limited by maximum depth, which was significantly highest in the control ponds (Fig. 6). The maximum depth was decreasing through time in all ponds, but more rapidly in the grazed and mowed ponds (Fig. S4). Reed had highest cover in the control ponds, while submergent macrophytes in the mowed ponds. Conductivity, pH, temperature and nutrient concentrations strongly fluctuated throughout the study period (Fig. S4), but did not differ between the exp. pond types except total phosphorus, which was lowest in the mowed ponds (Table 4, Fig. 6).

**Fig. 6.**
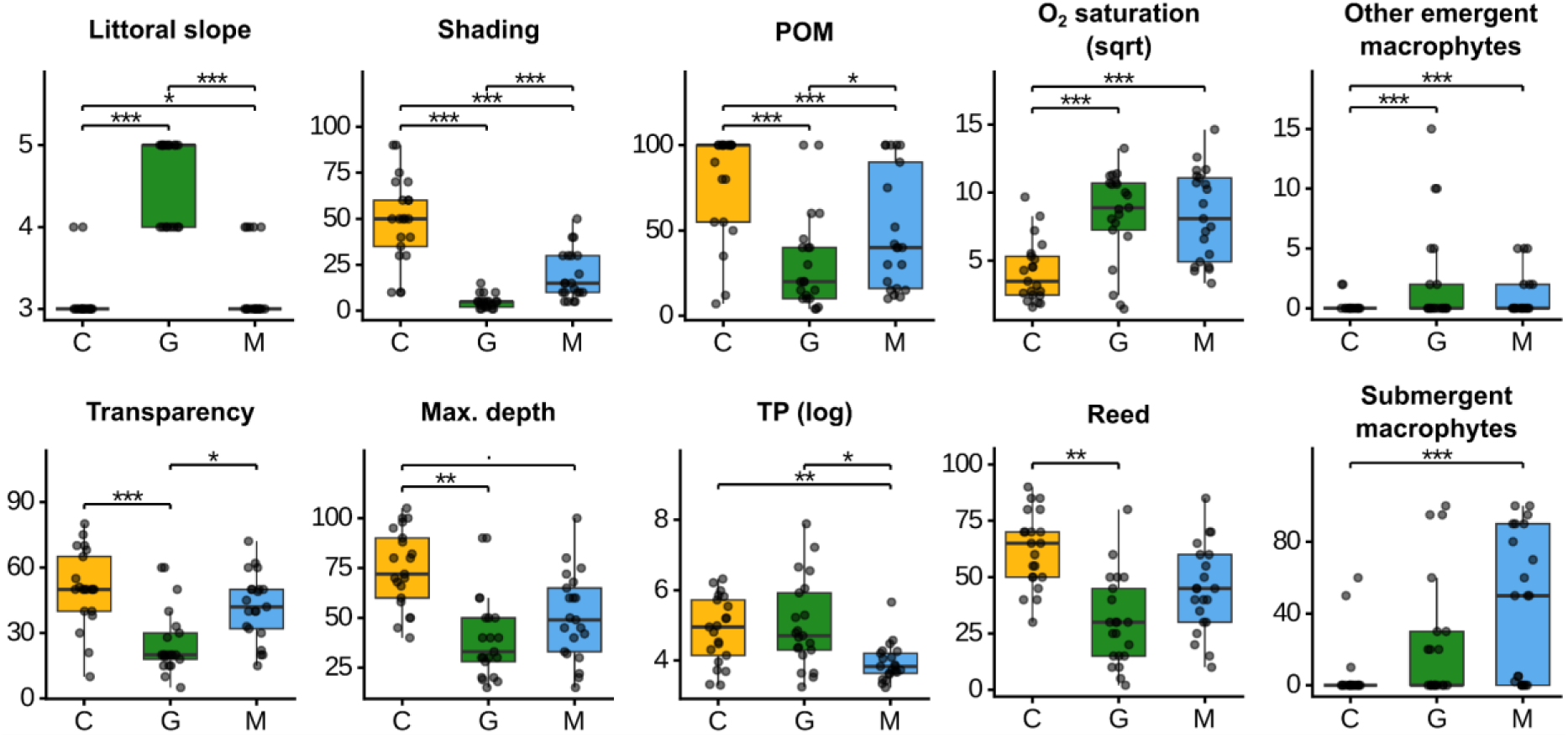
Boxplots of variables differing significantly between the types of experimental ponds (see Table 4; C – control, G – grazed, M – mowed). Boxes represent the median (horizontal line) and inter-quartile range (25^th^–75^th^ percentiles) and whiskers represent the range of values under 1.5 box-lengths. Points along boxplots stand for individual values. Signs above plots indicate significant differences between connected boxes. *** p < 0.001; ** p < 0.01; * p < 0.05; . p < 0.1.

**Table 4.**
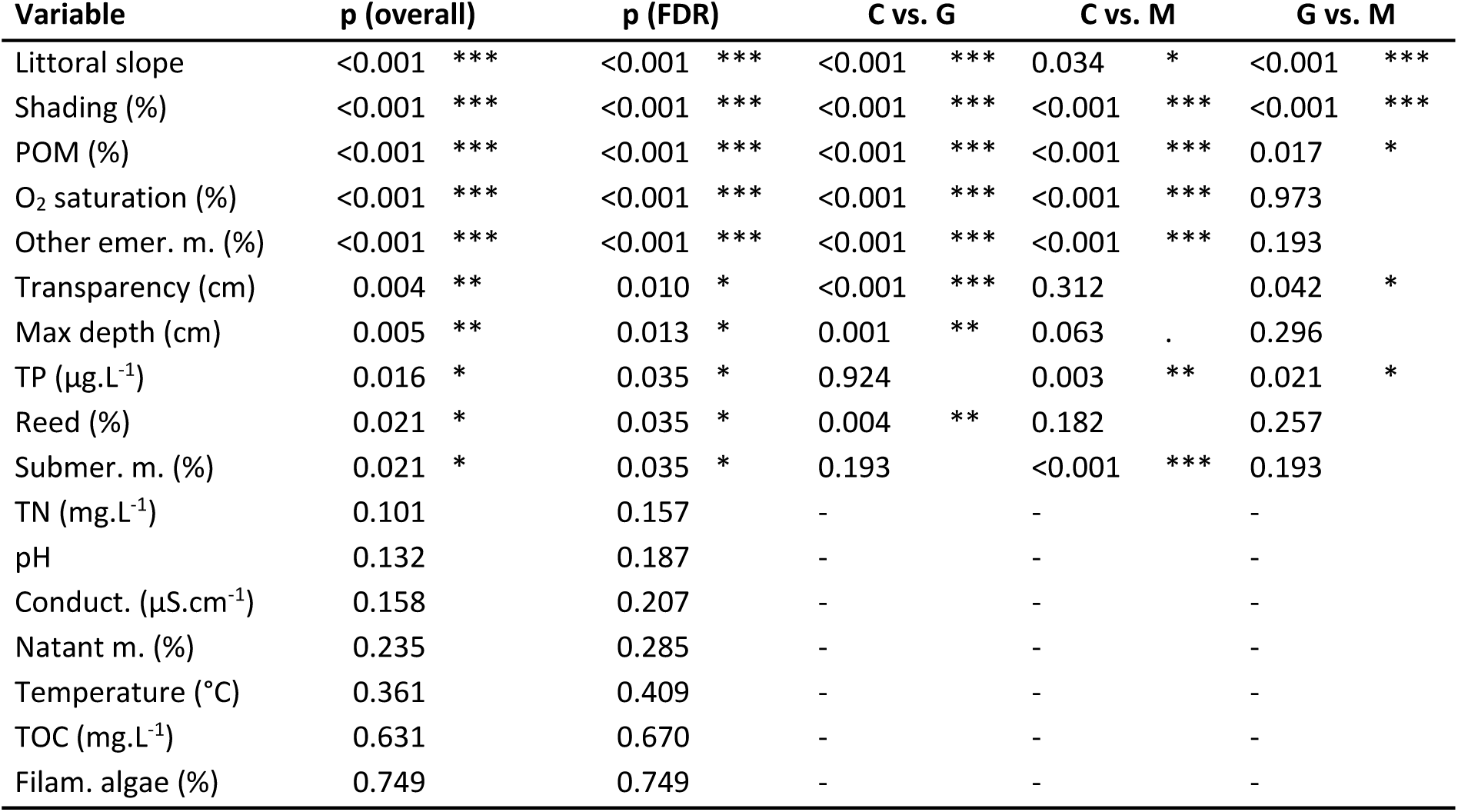
Summary of the nonparametric longitudinal model with post-hoc pairwise comparisons with FDR corrected p-values performed on variables measured in three types of experimental ponds. Data since June 2021 were used. Variables are ordered from the most significant to the least significant. For graphical visualisation of the results, see Fig. 6. p (overall) – uncorrected p-values of the model; p (FDR) – FDR corrected p-values of the model; C vs. G, C vs. M, G vs. M – FDR corrected p-values and significance from post-hoc pairwise comparisons of the experimental ponds (C – control, G – grazed, M – mowed). Other emer. m. – other emergent macrophytes; Submer. m. – submergent macrophytes; Conduct. – conductivity; Natant m. – natant macrophytes. *** p < 0.001; ** p < 0.01; * p < 0.05; . p < 0.1.

## 4. Discussion

### 4.1 Impact of management on macroinvertebrate diversity and habitat structure of the experimental ponds

Our experiment confirmed that the management regime had a significant effect on macroinvertebrate community composition and densities in the newly built ponds, but, interestingly, only a limited effect on species richness (Table 2, Fig. 3). We also detected a significant influence of the treatments on pond characteristics, mainly those describing habitat and vegetation structure (Table 4). The differences in species composition between treatments were relatively high throughout the entire study period, and generally increased over time (Fig. 5), as the management practices and succession gradually modified the ponds into three visibly distinct types of aquatic biotopes (Fig. S2). The ecology of indicator species identified for each treatment (Table 3) showed that the management effects on community structure and habitat conditions are tightly interconnected.

Both grazing and mowing reduced reed cover and height, which resulted in lower shading of the managed ponds compared with the control ponds. Grazing had a stronger effect than mowing, as the uninterrupted presence of livestock throughout the vegetation season ensured continuous biomass removal and trampling (Reeves & Champion 2004). Intensive reed reduction in the grazed ponds also enabled the growth of other emergent macrophytes not attractive to livestock (e.g. *Ranunculus sceleratus*). In contrast, shading of the mowed ponds tended to increase between mowing events due to rapid reed regrowth in the eutrophic environment. Nevertheless, the increased sunlight exposure in both types of managed ponds promoted the development of algae and submergent macrophytes, which was also reflected in higher water oxygen saturation. These favourable conditions likely improved food availability, which contributed to the higher macroinvertebrate densities observed in the managed ponds compared to the controls (Benke & Huryn 2010). The open character of the managed ponds attracted heliophilic taxa, particularly Odonata, which exhibited increased species richness and densities in the managed ponds. Odonata larvae were more abundant in the mowed than in the grazed ponds, possibly due to more stable submergent vegetation and higher water transparency, the latter being important for visually driven predators. All indicator taxa determined for the mowed ponds were macroinvertebrates dependent on dense submergent vegetation (the molluscs *Gyraulus crista* and *Lymnaea stagnalis*, the water bugs *Plea cryptica* and *Ilyocoris cimicoides*, and the dragonfly *Sympetrum striolatum*).

In the grazed ponds, livestock entering the water periodically decreased water transparency and physically disturbed submergent macrophytes. Grazing had a pronounced effect on the littoral zone morphology, as the livestock trampling reduced the littoral slope and overall depth of the grazed ponds (Table 4, Fig. 6). This resulted in the formation of a very heterogenous littoral zone, with emergence of extensive sun-exposed, non-vegetated patches covered by a thin film of water (Fig. S2A). Such heterogeneous microhabitat structure and extent of shallow littoral areas in ponds are known to support distinct communities of aquatic macroinvertebrates (Gleason et al. 2018, Oertli 2018). This was reflected in our study by the highest number of indicator taxa in the grazed ponds. These taxa include mainly beetles typically inhabiting shallow, warm, nutrient-rich ponds with muddy substrates and submergent vegetation (*Hygrotus impressopunctatus*, *Agabus bipustulatus*, *Helophorus* gr. *minutus*, *Enochrus quadripunctatus*). Other species are restricted to exposed muddy shorelines with accumulations of decaying plant debris (*Ochthebius bernhardi* and *O. minimus*) or with algal growths (mollusc *Radix labiata*). The remaining species are eurytopic taxa that prefer newly formed habitats (*Sigara nigrolineata*, *Hydroglyphus geminus*), and were likely frequent in the grazed ponds due to the livestock-generated disturbances maintaining microhabitats in early successional stages.

The control ponds were inhabited by taxa tolerant of low oxygen levels, high shading, dense reed cover and organic-matter rich substrates (Table 4, Fig. 6). Only two taxa were identified as indicators of the control ponds: mollusc *Aplexa hypnorum*, which inhabits a broad scale of vegetation-rich lowland waters including temporary pools and forested pond margins, and larvae of the beetle *Contacyphon* sp., which occur in accumulations of decaying plant material. These larvae are likely *C. lavipennis* (based on separate terrestrial macroinvertebrate surveys), a species inhabiting dense emergent littoral vegetation, typically reed stands.

The nutrient concentrations, pH and conductivity fluctuated considerably over time in all ponds, but did not significantly differ between the pond types. The exception were total phosphorus concentrations, which had lowest values in the mowed ponds. This phenomenon probably reflects their greatest distance from the nutrient-rich *Spálený potok* stream, resulting in the lowest hydrological connectivity (Tockner et al. 1999). Nevertheless, all studied ponds were extremely nutrient-enriched, with all measured values corresponding to eutrophic (TP values 24–96 μg.L^−1^) or hypertrophic class (TP > 96 μg.L^−1^; Carlson & Simpson 1996). Conductivity was also high (values between 1000 and 4400 μS.L^−1^), reflecting the historical salt-marsh character of the wetland (Kotasová Adámková et al. 2022). These conditions possibly limited the potential macroinvertebrate assemblage of the experimental ponds only to taxa adapted to such specific environmental conditions. The comparable water quality across treatments supports our conclusion that the observed differences in invertebrate communities were driven primarily by divergent habitat and vegetation structure.

### 4.2 Temporal trends of macroinvertebrate communities of the experimental ponds and whole wetland context

Our observations indicate that newly created ponds can be rapidly colonised by actively flying macroinvertebrates, consistent with previous studies (e.g. Williams et al. 2008). Two weeks after excavation, the experimental ponds were already inhabited by a small number of Heteroptera and Coleoptera species. This initial community consisted of pioneer species of water boatmen (Corixidae) and backswimmers (Notonectidae), as well as eurytopic beetles inhabiting broad scale of biotopes (e.g. *Agabus bipustulatus*, *Hygrotus inaequalis*, *Rhantus suturalis* and *R. frontalis*). Both species richness and densities increased over the following year and a half, eventually reaching a peak. We assume this pattern reflected the specific conditions of the early successional stage, characterised by low cover of emergent macrophytes and limited organic matter, but already supporting developed submergent vegetation, algae and plankton. As a result, pioneer taxa, eurytopic taxa and vegetation-associated taxa co-occurred during this period. This peak was followed by complete drying of most experimental ponds during the summer 2022, after which only a few taxa and individuals were recorded in the next sampling. Since then, both richness and densities have partially recovered in most ponds, although the values have not returned to pre-drought levels. Based on the observed trends, predicting future community development in the experimental ponds remains difficult. The hydrological regime of the wetland, involving frequent flooding and periodic drought, may promote an oscillatory dynamic in which succession is periodically reset by these events (Ruhí et al. 2016, O’Neill 2016).

We suggest that the creation of small experimental ponds increased the overall biodiversity of the wetland. Their macroinvertebrate communities differed not only between management treatments, but also among individual ponds within treatments and across sampling periods, which contributed to high beta and gamma diversity. The experimental ponds provided habitat for many macroinvertebrates that were absent or only sparsely recorded in the old pond (Table S1). These included taxa associated with early-successional habitats or with small and temporary standing waters – common pioneer taxa as well as species rare in Czechia (e.g. beetles *Berosus frontifoveatus* and *Bidessus nasutus*; Hejda et al. 2017). On the other hand, few taxa requiring larger or permanent aquatic biotopes usually with dense submergent vegetation were found only in the old pond (e.g. the beetles *Hydrophilus piceus*, *Ilybius fenestratus* and *I. subaeneus*, and the damselfly *Erythromma viridulum*). This pattern indicates that a mosaic of aquatic biotopes differing in size, successional stage and water-level stability contributes substantially to the preservation of local biodiversity (Sinclair et al. 2021, Fahy et al. 2025).

### 4.3 Implications for artificial pond management

Our results have several implications for the management of newly constructed ponds in eutrophic landscapes. These artificial biotopes are often adversely affected by rapid expansive vegetation proliferation (Fluet-Chouinard et al. 2023) and, in many cases, by inappropriate design resulting in low littoral heterogeneity (Zamora-Marín et al. 2021, Fahy et al. 2025). We suggest that both issues can be mitigated through appropriate long-term grazing or mowing. Both management regimes suppressed expansive vegetation, increased sunlight exposure, and enhanced habitat availability for macroinvertebrates in the experimental ponds. Moreover, each regime supported specialised taxa and influenced habitat structure in an unique way, indicating that combining multiple management approaches within a single area may increase the overall diversity of wetland pondscapes. Both grazing and mowing are traditional ways of wetland use in Europe, providing a precedent for their application as restoration tools (van Diggelen et al. 2006, Biró et al. 2019). However, several lines of evidence suggest that grazing may be a superior management regime compared with mowing. As noted above, grazing continuously removes vegetation cover, maintaining consistently high light availability, whereas mowed ponds alternate between periods of reduced and increased sun exposure. Furthermore, although our experimental ponds were initially excavated as simple bowl shapes, livestock trampling gradually created heterogenous littoral zones with smooth transition between aquatic and terrestrial habitats. In effect, this process produced the desired pond morphology at negligible additional costs, eliminating the need for expensive engineering of complex pond shapes. Livestock trampling can also generate small pools and ephemeral water-filled depressions, further increasing habitat complexity and fine-scale heterogeneity (Biró et al. 2019). Although our study focused on very small ponds (approximately ∼20 m^2^ in area and less than 1 m deep), grazing may also plausibly benefit the littoral zones of larger ponds by increasing light availability and promoting structurally diverse mosaic of microhabitats (Middleton 2016, Fløjgaard et al. 2022). In larger wetland complexes containing numerous ponds of varying morphology, grazing alone may generate substantial between-pond heterogeneity, as livestock tend to prefer certain ponds over others (Morris & Reich 2013).

Studies examining the effects of wetland grazing on aquatic invertebrate diversity have yielded different results. While some studies report positive impacts (Marty 2005, Davis & Bidwell, 2008), others are more equivocal (Steinman et al. 2003), and intensive grazing of wetlands has been shown to negatively affect the macroinvertebrate communities (Bloechl et al. 2010, Epele & Miserendino 2015). In our extensively grazed wetland, species richness in grazed ponds was not significantly different from other types, but neither did we observe any deleterious effects. Instead, grazed ponds exhibited the highest macroinvertebrate densities and provided habitat for highly specialised taxa. Importantly, although the nominal stocking rate of 2 livestock units per hectare would typically be considered intensive (AOPK ČR 2021), the functional effects in our eutrophic system corresponded to extensive grazing. High primary productivity of the pasture buffered grazing pressure, preventing habitat degradation and maintaining structural heterogeneity. These findings suggest that grazing intensity should be interpreted relative to site-specific productivity rather than inferred solely from livestock density.

A pervasive issue in artificial pond construction is colonisation by non-native fish, which can severely degrade habitat quality (Janáč et al. 2025). In Czechia, this is exacerbated by the tendency to construct large, permanent ponds (Sychra et al. 2025). On the other hand, small and shallow ponds are often overlooked, despite the clear benefits of their susceptibility to drying. Periodic drying can prevent the establishment of permanent fish stock (Moor et al. 2024, Janáč et al. 2025) and we suspect that drying of our experimental ponds helped prevent fish colonisation. Additionally, drying resets communities to early successional stages, increasing temporal species turnover and creating opportunities for pioneer taxa and temporary-water specialists (Collinson et al. 1995, Ruhí et al. 2016, O’Neill 2016), which were frequently observed in the experimental ponds throughout the entire study.

Since our study focused on a single locality, it should be viewed as a field experiment illustrating the potential of different management regimes. Nevertheless, the results are robust and offer practical guidance for improving biodiversity in artificial ponds and we believe these findings are highly transferable to other similar eutrophic environments. It is important to acknowledge that additional factors may limit restoration potential of the studied area. Firstly, beyond high nutrient loads, we cannot exclude the presence of other pollutants not measured in this study, including heavy metals and pesticides, both known to be toxic to aquatic fauna (Schulz et al. 2021, Kadiru et al. 2022, Jeong et al. 2023). Secondly, due to significant degradation and loss of most Czech lowland wetlands (Skaloš et al. 2017), the regional macroinvertebrate species pool is likely depleted, potentially limiting colonisation of newly constructed ponds. Despite these constraints, the introduction of management regimes had clear positive effects on the wetland. Our study demonstrates that grazing and mowing can induce rapid and substantial ecological changes even in an eutrophic environment and hold strong potential for enhancing aquatic biodiversity. Further research, particularly across multiple managed wetland pondscapes, is needed.

### 4.4 Concluding remarks

Taken together, our results suggest that the ecological effects of grazing and mowing in eutrophic wetland pondscapes are shaped less by nominal management intensity and more by the interaction between biomass production, disturbance regime, and habitat structure. Even in nutrient-rich agricultural landscapes with a potentially depleted regional species pool, appropriately adjusted management can maintain structural heterogeneity of artificial ponds, facilitate colonisation, and support specialised taxa. Rather than focusing solely on pond creation, restoration strategies should prioritise long-term management regimes that sustain open, dynamic and spatially heterogeneous aquatic habitats. In highly modified lowland landscapes, such management may represent one of the most realistic and cost-effective approaches for conserving aquatic biodiversity.

## Acknowledgements

We are grateful to all our colleagues who participated in the fieldwork, especially Stanislav Němejc, Marcela Růžičková, Alžbeta Devánová, Barbora Řehůřková and many others. We would like to thank Erika Šlachová for the determination of the mollusks. The study was supported by the project “Land management and assessment of its impact on wetland biodiversity” (TJ04000145) by the Technology Agency of the Czech Republic.

## Author contributions

Conceptualization: Jana Petruželová, Marie Kotasová Adámková, Jan Petružela; Data curation: Jana Petruželová, Alexandra Černá; Formal analysis: Jan Petružela, Jana Petruželová; Funding acquisition: Marie Kotasová Adámková; Field Work and Investigation: Jana Petruželová, Jan Petružela, Alexandra Černá, Marie Kotasová Adámková; Methodology: Jana Petruželová, Jan Petružela, Marie Kotasová Adámková; Project administration: Marie Kotasová Adámková; Supervision: Marie Kotasová Adámková; Writing – original draft: Jan Petružela, Jana Petruželová; Writing – review and editing: Jana Petruželová, Jan Petružela, Alexandra Černá, Marie Kotasová Adámková

## Data Statement

All data used in this study is available at the Mendeley Data repository under DOI: 10.17632/cdvv72kn3p.1 (Petruželová et al. 2026)

## Notes

### Competing Interest Statement

The authors have declared no competing interest.

