## Supplementary Figures S1-S4 for "Grazing and mowing enhance aquatic macroinvertebrate diversity of small artificial ponds in eutrophic landscape"

[*] shared first authorship

[+] corresponding author

Fig. S1. Illustrative photos of the experimental wetland. (A) The wetland in May 2020, before the management practices were implemented. The locations of the managed and unmanaged areas are indicated in the picture (see also Fig. 1; photo acquired using an unmanned quadcopter registration number OK-X071A by L. Tichý). (B) The old pond in June 2024 (photo: J. Petruželová). (C) Mowed area in May 2021 (photo: J. Petruželová). (D) Grazed area in April 2024 (photo: M. Kotasová Adámková).

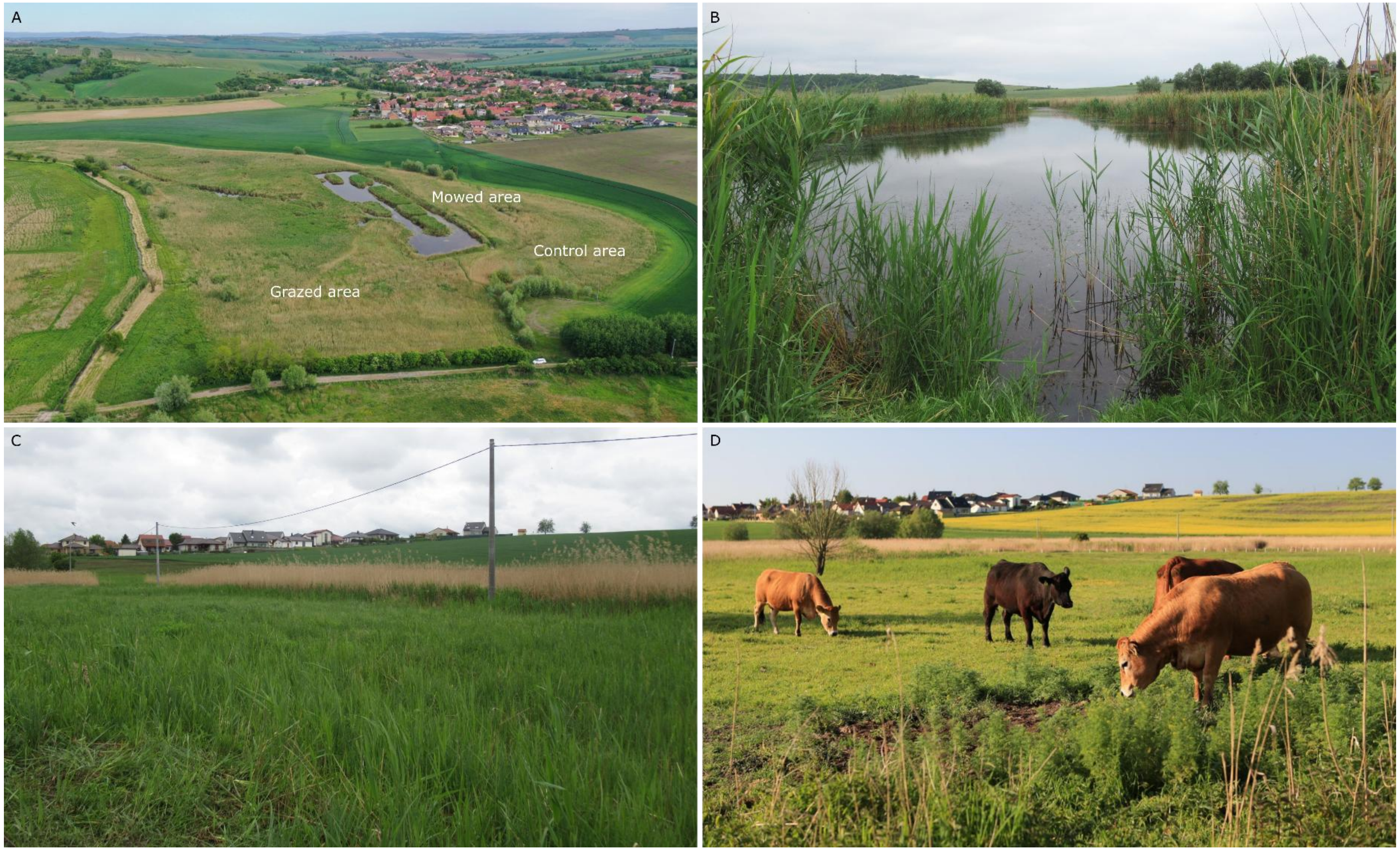

Fig. S2. Photos of selected experimental ponds from all sampling periods: A – Grazed pond (G2); B – Mowed pond (M1); C – Control pond (C1). Photos: J. Petruželová, J. Petružela, M. Kotasová Adámková.

A: Grazed pond – G2

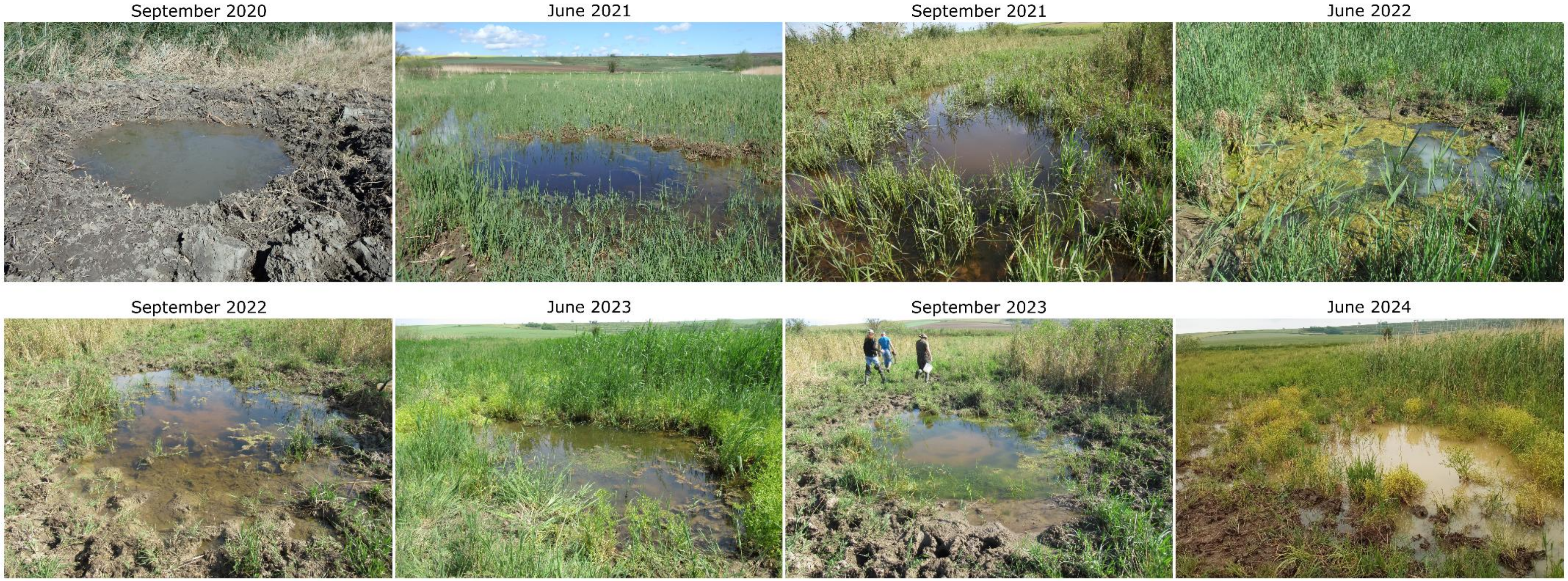

B: Mowed pond – M1

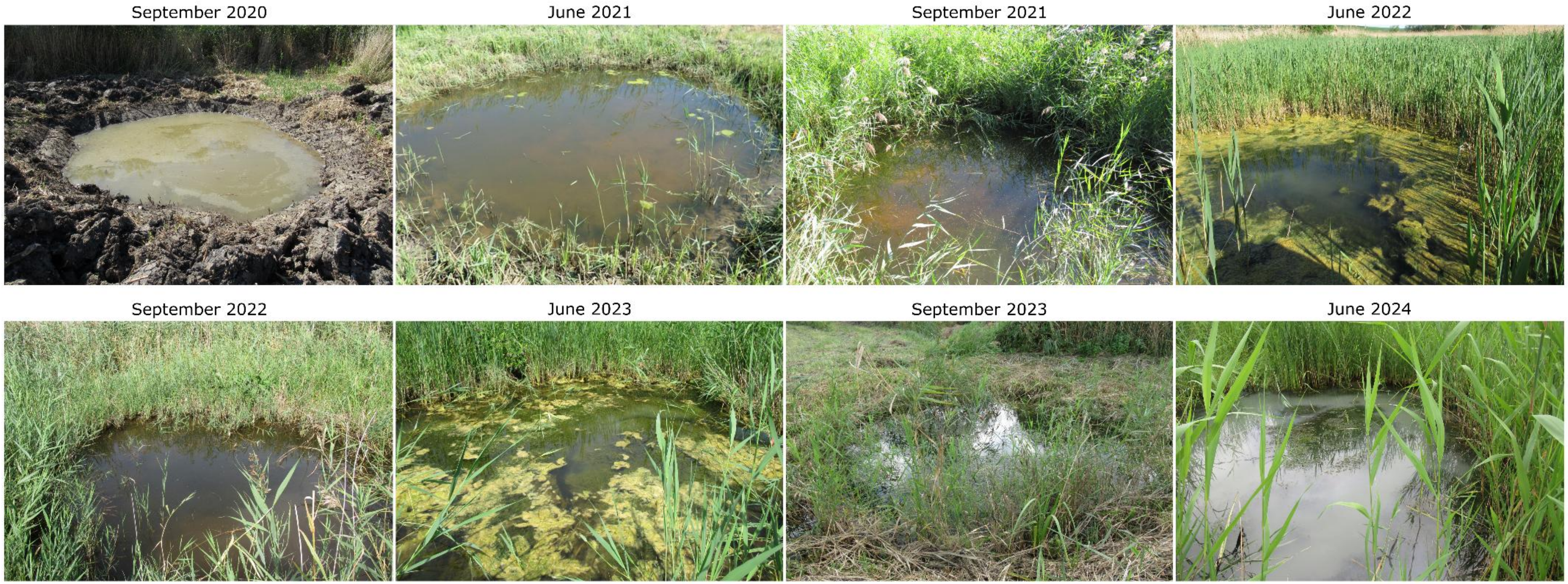

C: Control pond – C1

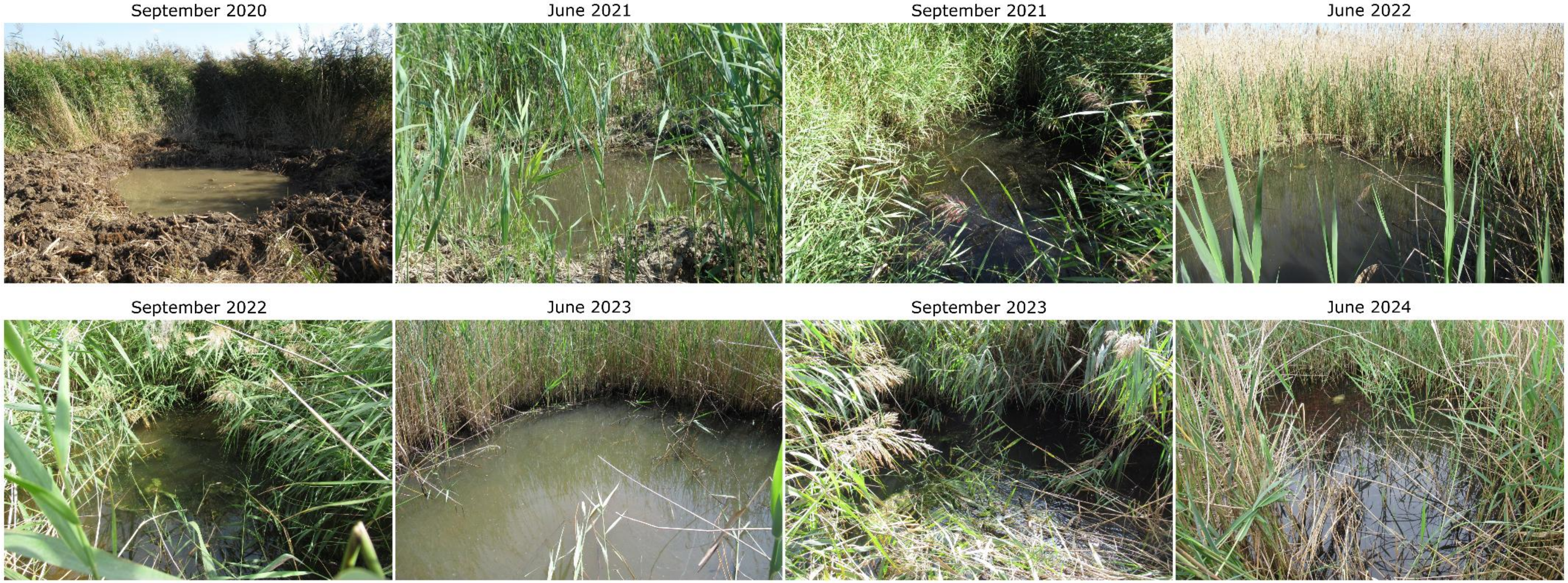

Fig. S3. Species richness and density of individual groups of macroinvertebrates at each sampling time and aquatic biotope. X axis stands for sampling terms between June 2020 and June 2024 (J – June, S – September, 20 – 2020, 21 – 2021, 22 – 2022, 23 – 2023, 24 – 2024).

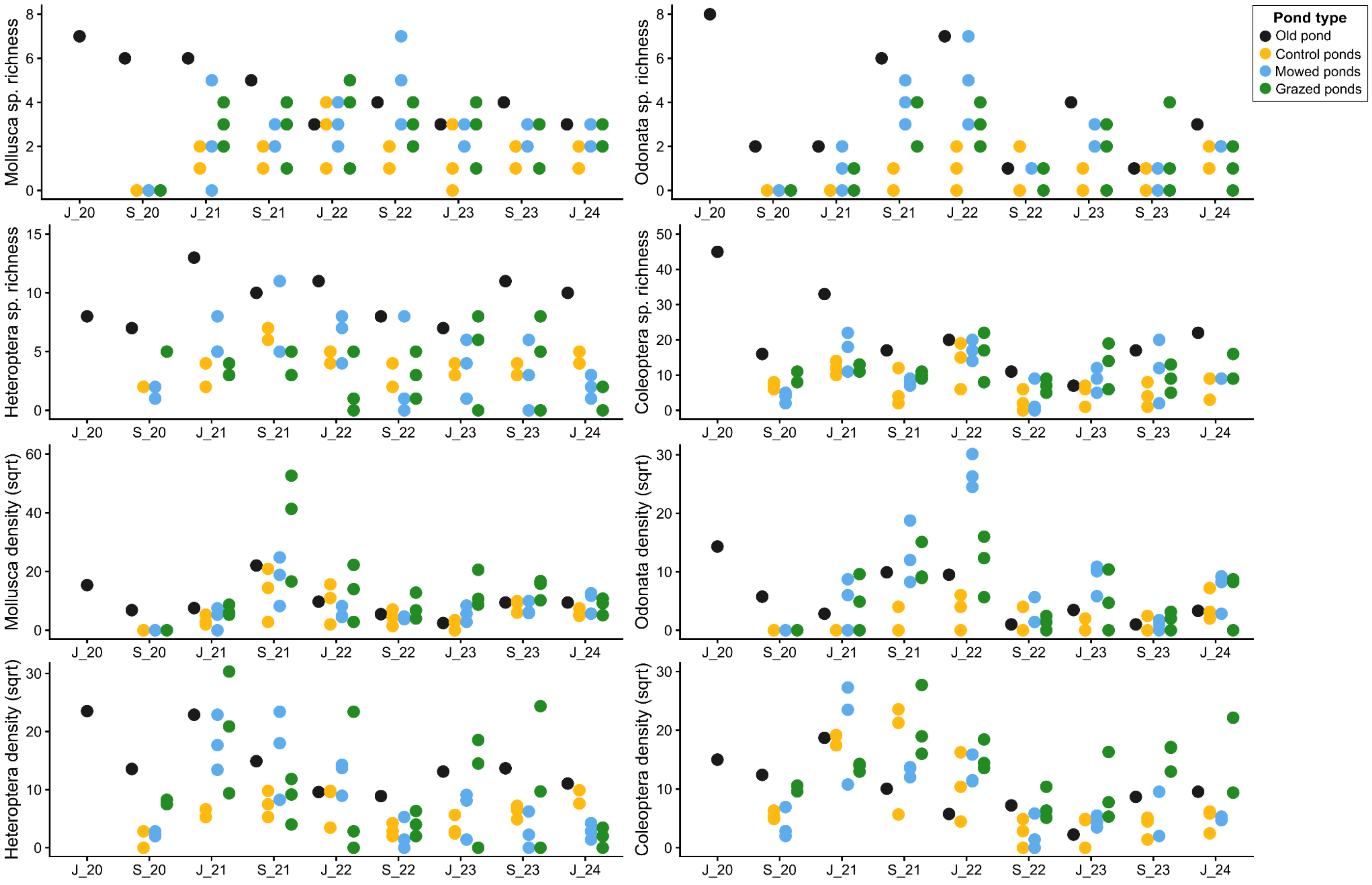

Fig. S4. Physico-chemical and habitat variables development through the study period at individual ponds. For measured units of the variables see Table 4. X axis stands for sampling terms between June 2020 and June 2024 (J – June, S – September, 20 – 2020, 21 – 2021, 22 – 2022, 23 – 2023, 24 – 2024).

A

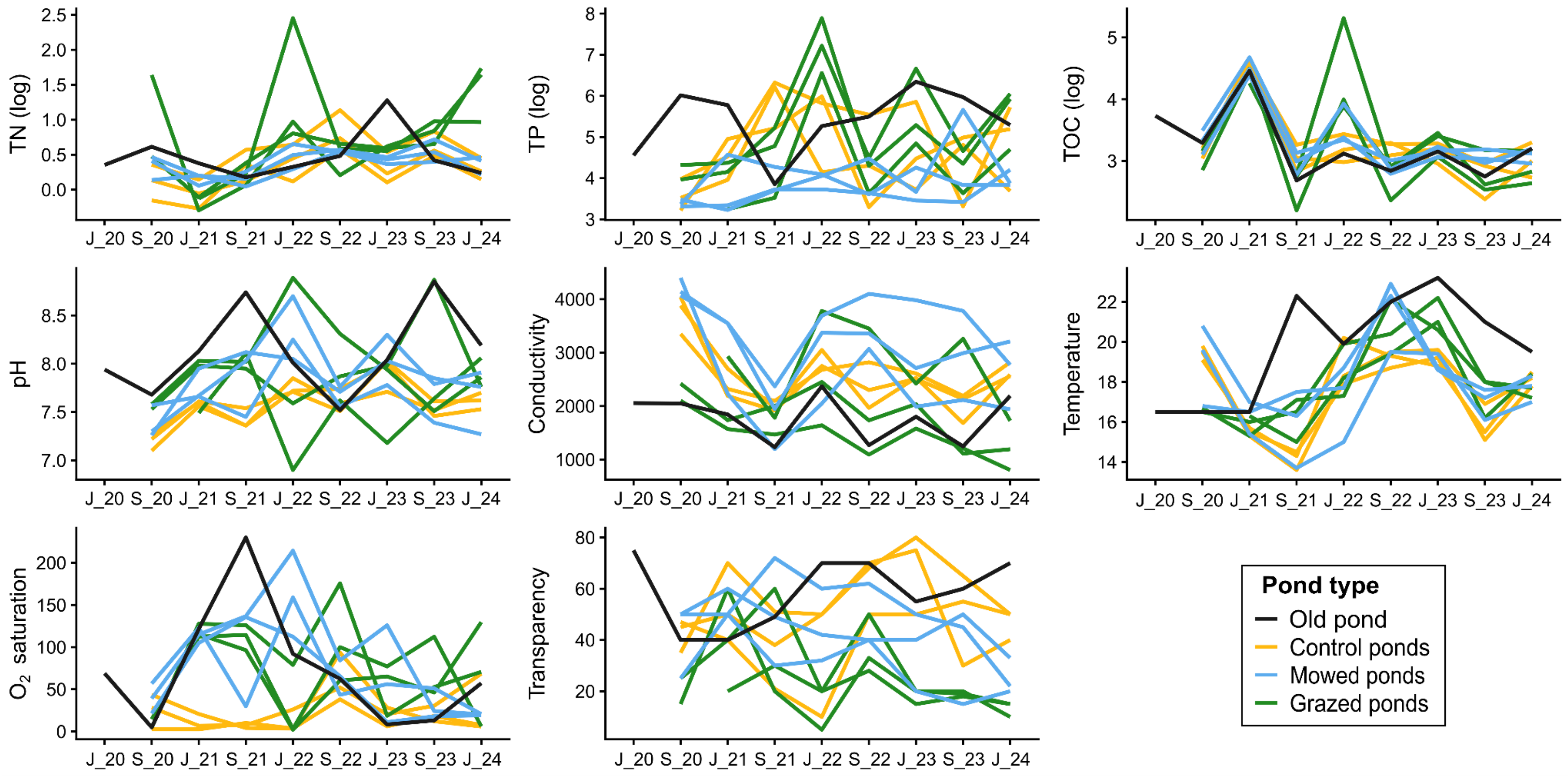

B

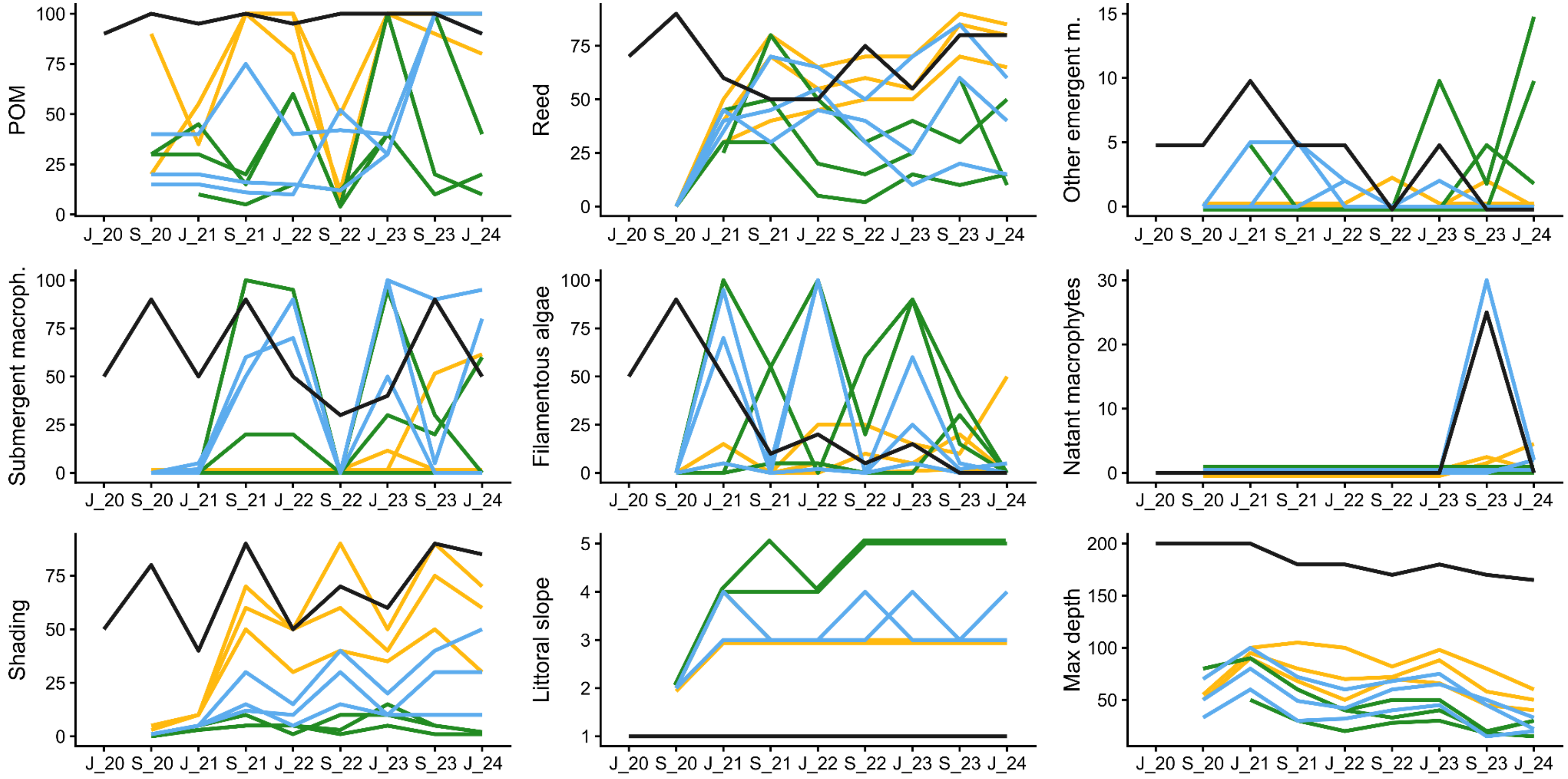

POM
Reed
Other emergent m.
Submergent macroph.
Filamentous algae
Natant macrophytes
Shading
Littoral slope
Max depth
J_20 S_20 J_21 S_21 J_22 S_22 J_23 S_23 J_24
